# More than just hitchhikers: a survey of bacterial communities associated with diatoms originating from marine reptiles

**DOI:** 10.1101/2022.04.19.488760

**Authors:** Klara Filek, Liesbeth Lebbe, Anne Willems, Peter Chaerle, Wim Vyverman, Marta Žižek, Sunčica Bosak

## Abstract

Diatoms and bacteria are known for being the first colonizers of submerged surfaces including the skin of marine reptiles. Sea turtle carapace and skin harbour diverse prokaryotic and eukaryotic microbial taxa, including several epizoic diatoms. However, the importance of diatom-bacteria associations is hardly investigated in biofilms associated with animal hosts. This study provides a detailed inventory of diatoms, bacteria, and diatom-associated bacteria originating from several loggerhead sea turtles using a combination of metabarcoding and culturing approaches. Carapace and skin samples *rbcL* and 16S rRNA amplicon sequencing showed a high diversity of diatoms and bacteria, respectively. Cultures of putative epizoic and non-epizoic diatom strains contained from 18 to 101 bacterial amplicon sequence variants (ASVs) and their bacterial assemblages strongly reflected those of their source host. Diatom strains allowed for enrichment and isolation of rare-in-source bacterial families such as *Marinobacteraceae, Alteromonadaceae*, and *Alcanivoracaceae*. When accounting for phylogenetic relationships between bacterial ASVs, we observed related diatom genera might retain related microbial taxa in culture, regardless of the source environment. These data provide deeper insights into several levels of sea turtle epizoic diatom and bacterial communities, and reveal the potential of epizoic biofilms as a source of novel microbes and possibly important diatom-bacteria associations.

## Introduction

Bacterial close interactions with microbial eukaryotes (protists) are common across different environments including marine habitats (Husnik *et al*. 2021). Despite the extent of genomic and metabolic diversity of microbial eukaryotes, and their importance in biogeochemical cycles, most of the information on host-microbe associations has been acquired from studies on animal hosts, particularly the digestive system of mammals (Thompson *et al*. 2017; Husnik *et al*. 2021). However, bacterial associations with microbial eukaryotes have been increasingly studied in ciliates, amoeba, dinoflagellates, and diatoms, traditionally in terms of endosymbiosis (as plastids or housed within the host cytoplasm, nucleus, or mitochondria) and as ectosymbionts (microalgal phycosphere). The range of host-bacteria interactions span from beneficial, commensal, to harmful (e.g., B_12_ vitamin production by bacteria, utilization of host-derived organic matter, competition for resources, antimicrobial compounds production by hosts) sometimes even expanding the hosts metabolic “toolbox”, but the types of interactions are often overlapping and difficult to decouple (Amin, Parker and Armbrust 2012; Seymour *et al*. 2017; Henry *et al*. 2021; Husnik *et al*. 2021; Boscaro *et al*. 2022).

Diatoms (Bacillariophyceae) are essential and omnipresent primary producers in aquatic environments, responsible for approximately 20% of oxygen production and 40% of primary production and particulate carbon export (Field *et al*. 1998; Jin *et al*. 2006; Tréguer *et al*. 2018). The bulk of research on planktonic diatoms in the water column and benthic diatoms inhabiting sediment biofilms revealed the importance of diatom-bacteria interactions (Amin, Parker and Armbrust 2012) and bacterial influence on diatoms’ community composition and productivity (Koedooder *et al*. 2019), growth and cell division (Amin *et al*. 2015; van Tol, Amin and Armbrust 2017), and sexual reproduction (Cirri, Vyverman and Pohnert 2018; Cirri *et al*. 2019), while diatoms can directly modulate the bacterial community via secondary metabolite production (Fei *et al*. 2020; Shibl *et al*. 2020). Efforts in elucidating the diatom-bacteria associations and interactions are still restricted to somewhat familiar systems of laboratory cultures, plankton, or sediment, while studies that expand the diatom-bacteria associations repertoire in other habitats remain scarce.

Marine vertebrates have been reported to be extensively colonized by diatoms along with other macro- and microorganisms as reported by morphology-based approaches and metabarcoding (Frick and Pfaller 2013; Rivera *et al*. 2018; Hooper *et al*. 2019; Blasi *et al*. 2021; Robinson and Pfaller 2022; Kanjer *et al*. 2022 preprint). There are multiple reports on novel diatom taxa associated with sea turtles and their potentially exclusive epizoic lifestyle as they have not yet been found elsewhere (Majewska *et al*. 2015, 2017, 2020; Riaux-Gobin *et al*. 2021). Microbial diversity observed on marine vertebrate epidermis suggests marine animals could be “hot spots” for microbial diversity and interactions in often nutrient-poor open seas (Keller *et al*. 2021). In this study we focused on surface microbial communities of loggerhead sea turtles in the Adriatic Sea. Reports on prokaryotic and microeukaryotic data on loggerhead sea turtles show diatoms make up a noticeable portion (up to 25%) of microbial eukaryotes on carapace and skin of loggerheads, often within complex biofilms dominated by bacterial phyla Proteobacteria (classes Gammaproteobacteria and Alphaproteobacteria), Bacteroidota, Bdellovibrionota, Cyanobacteria, and Firmicutes (Blasi *et al*. 2021; Kanjer *et al*. 2022 preprint). It is still unknown if sea turtle-associated microbial epizoic communities have any effect on their host or *vice versa*, including putative epizoic diatoms. Nonetheless, it is becoming clear loggerhead sea turtle carapace and skin are dynamic and microbially-rich environments with a potential to act as a reservoir of diverse and novel microbial species (Kanjer *et al*. 2022 preprint). Insights into the biodiversity of marine vertebrate host-derived diatoms and diatom-associated bacteria are still lacking, even though they could be crucial for understanding the biology and lifestyle of epizoic diatoms.

The aim of this study was to provide an inventory of diatom, bacterial, and diatom-associated bacterial communities originating from several loggerhead sea turtles via marker gene microbial profiling (*rbcL* and 16S rRNA genes) and cultivation. The main objectives of this study were (i) to examine the microbial community structure on the surface of loggerhead sea turtles, (ii) to isolate and cultivate turtle-associated diatom strains, (iii) to profile the bacterial communities associated with individual diatom strains in culture, and (iv) to isolate and cultivate bacteria from several diatom strains to sample the culturable range of diatom-associated bacterial communities. This multi-layered approach provides a deeper understanding of sea turtle epizoic biofilm potential as a source of novel microbes, source-to-culture bacterial shifts in diatom strains, and potentially important diatom-bacteria associations in epizoic biofilms.

## Materials and methods

### Loggerhead carapace and skin sampling

The samples in this study were collected from four loggerhead sea turtles in the Adriatic Sea during 2019. Living samples of carapace (randomised collection across the whole surface) and skin (head, neck, and flippers) biofilms used for diatom cell isolations were collected by sterile toothbrushes and resuspended in 50 ml conical sterile tubes containing filtered sea water (Table 1). Samples intended for microbial metabarcoding were collected as described above and resuspended in 50 ml sterile conical tubes containing 96% ethanol for preservation at −20 °C until further processing (Kanjer *et al*. 2022 preprint). Live samples intended for diatom isolation were diluted in sterile f/2 culture medium (Sigma-Aldrich, Germany) (Guillard’s medium for diatoms; Guillard 1975) with salinity matching the sample collection source, either in sterile petri dishes (LLG, Germany) or flat bottom transparent 6 or 24 well plates (Guangzhou Jet Bio-Filtration, China), and incubated at 18-20 °C at 7-10 μmol m^2^ s^-1^, 12:12 (light:dark) cycle.

**Table 1.** Description of diatom strains used in this study. Turtle ID includes carapace and skin source samples taken from a single turtle.

| Diatom strain ID | Turtle ID | Source sample (ID) | Scientific name | Host Relation** | Time from isolation to harvest (days) | Bacterial isolates |
| --- | --- | --- | --- | --- | --- | --- |
| DM0150 | ID010 | Carapace (TB139) | *Achnanthes squaliformis* | epizoic | 388 | yes |
| DM0177 | ID010 | Carapace (TB139) | *Achnanthes squaliformis* | epizoic | 332 | yes |
| DM0178 | ID010 | Skin (TB140) | *Nitzschia* sp. | non-epizoic | 343 | no |
| DM0179 | ID010 | Skin (TB140) | *Entomoneis* sp. | non-epizoic | 332 | no |
| DM0052 | ID034 | Carapace (TB89) | *Achnanthes elongata* | epizoic | 667 | no |
| DM0053 | ID034 | Carapace (TB89) | *Achnanthes elongata* | epizoic | 682 | yes |
| DM0054 | ID034 | Carapace (TB89) | *Achnanthes elongata* | epizoic | 673 | no |
| DM0060 | ID034 | Carapace (TB89) | *Achnanthes elongata* | epizoic | 647 | yes |
| DM0070 | ID034 | Carapace (TB89) | *Amphora* sp. 1 | non-epizoic | 649 | no |
| DM0077 | ID034 | Skin (TB90) | *Poulinea lepidochelicola* | epizoic | 644 | yes |
| DM0123 | ID047 | Carapace (TB115) | *Diploneis* sp. | non-epizoic | 503 | no |
| DM0129 | ID047 | Carapace (TB115) | *Diploneis* sp. | non-epizoic | 507 | yes |
| DM0136 | ID047 | Carapace (TB115*) | *Diploneis* sp. | non-epizoic | 512 | no |
| DM0147 | ID047 | Carapace (TB115*) | *Fallacia* sp. | non-epizoic | 495 | yes |
| DM0168 | ID073 | Carapace (TB155) | *Achnanthes elongata* | epizoic | 365 | no |
| DM0170 | ID073 | Carapace (TB155) | *Achnanthes elongata* | epizoic | 367 | yes |
| DM0181 | ID073 | Carapace (TB155) | *Psammodictyon panduriforme* | non-epizoic | 340 | yes |
| DM0182 | ID073 | Carapace (TB155) | *Amphora* sp. 2 | non-epizoic | 356 | no |
| DM0183 | ID073 | Carapace (TB155) | *Psammodictyon* sp. | non-epizoic | 333 | yes |

\*In case of ID047 a second carapace sample was obtained a month after the initial one (TB115) and it did not undergo NGS sequencing.
\*\*Host relation status (epizoic or non-epizoic) is based on Ashworth *et al.* (2022; preprint at https://doi.org/10.21203/rs.3.rs-1041030/v2)

### Diatom isolation, culturing, and identification

#### Establishing monoclonal diatom cultures

Diatoms were isolated from the diluted carapace and skin source samples within the next 8-10 weeks by weekly screenings using an inverted light microscope (Olympus CKX41, Olympus, Tokyo, Japan) and manual isolation of single diatom cells by micropipetting (Andersen 2005). Monoclonal xenic cultures were established by passaging a single cell through multiple series of sterile f/2 medium which facilitated removal of visible eukaryotic contaminants and preservation of bacteria in the diatom phycosphere (Andersen 2005). For the remainder of the study, diatom strains were grown in 25 cm^2^ or 75 cm^2^ cell culture flasks (VWR Avantor, USA), in 17 ml or 50 ml f/2 medium, respectively, at conditions as described above. Upon reaching higher densities (late exponential or stationary phase) the strains were sub-cultured. The cultures were subsampled for morphological and molecular analyses upon reaching late exponential phase by pelleting and removing the excess culture medium. Pellets for morphological analyses were stored at 4 °C in at least 70% EtOH, while pellets for marker gene analysis were stored at −20 °C in 96% EtOH. All diatom strains in this study (Table 1) are available at the BCCM/DCG culture collection (https://bccm.belspo.be/about-us/bccm-dcg; see Supplementary Table S1 for extended metadata and S2 for culture collection codes).

#### Diatom identification via morphology and *rbcL* sequencing

Diatom pellets were cleaned of organic matter from silicate frustules following Simonsen’s cleaning method (Simonsen 1974; Hasle 1978). Diatom samples (5 ml volume in ethanol) were washed with distilled water prior to adding an equal volume of saturated KMnO_4_ solution and incubating for 24 hours at room temperature. After 24 hours an equal volume of concentrated HCl was added to the samples, which were heated over an alcohol burner flame, and then washed with distilled water until neutral pH (approximately five times). Permanent slides were prepared by drying cleaned frustules on 22×22 mm coverslips (Hirschmann, Germany) and mounting with Naphrax (Brunel Microscopes Ltd, Chippenham, UK). Permanent slides were analysed with Zeiss Axio Imager A2 with DIC and an Axiocam 305 digital camera (Carl Zeiss, Jena, Germany). Stubs for scanning electron microscopy (SEM) analyses were prepared by drying cleaned frustules onto 3 μm pore size (13 mm in diameter) Nucleopore polycarbonate membrane filters (Pleasanton, CA, USA) before sputter-coating. Dried filters were mounted on aluminium stubs with carbon tape and sputter-coated with platinum (10 nm) using a Precision Etching and Coating System, PECS II (Gatan Inc., Pleasanton, CA, USA). The specimens were analysed with JEOL JSM-7800F scanning electron microscope (JEOL, Tokyo, Japan).

For molecular identification via the *rbcL* gene, diatom DNA was extracted from the pellets by DNeasy Plant Mini Kit (Qiagen) following the manufacturer’s instructions with an extra pre-processing bead beating step for disrupting the diatom colonies and frustules. The pellets were mixed with 0.5 g of 1.0 mm glass beads (Qiagen) in a sterile 2 ml safe lock microtube and vortexed horizontally at maximum speed for 10 minutes prior to continuing with the manufacturer’s protocol. The extracted DNA was used as a template to amplify the *rbcL* marker as described in Theriot *et al*. (2015). The initial PCR reactions were performed in 25 μl volume reactions consisting of 1 μl of DNA template, 12.5 μl Takara EmeraldAmp Master Mix 2x (Takara Bio, Japan), 0.5 μl primers rbcL40+ and rbcL1444- (0.2 μM final concentration), 10.5 μl of sterile dH_2_O, while nested PCR reactions were performed in 50 μl (double the reagents for 25 μl reaction) with similar set up as above but different reverse primer rbcL1255- (see Supplementary Table S2 for primers list). The thermocycling conditions for the initial sreaction were 94 °C for 5 min, 30 cycles of 98 °C for 10 sec, T_a_^(initial)^ = 50 °C or T_a_^(nested)^ = 50 °C for 60 sec, 72 °C for 2 min, and final extension at 72 °C for 15 min. The amplicons were purified by the NucleoSpin Gel and PCR cleanup kit according to manufacturer’s instructions (Macherey-Nagel, Dueren, Germany). Purified products were sent for Sanger sequencing with primers rbcL404+ and rbcL587-to Macrogen (http://dna.macrogen-europe.com).

Diatom strains identified as *Achnanthes elongata, Achnanthes squaliformis*, and *Poulinea lepidochelicola* were considered to be exclusively epizoic diatoms, while other diatoms were categorized as non-epizoic for the purposes of this study; based on Ashworth *et al*. (2022, preprint).

### Microbial community profiling in source samples and diatom strains

#### Source samples processing and amplicon sequencing

Carapace and skin samples’ (preserved in ethanol) DNA was extracted from approximately 250 mg of the sample pellet in duplicates using the DNeasy PowerSoil kit (Qiagen) following the manufacturer’s guidelines with several modifications as described in Kanjer *et al*. (2022, preprint): PowerBead tubes with samples were incubated in a sonicator at 50 °C at 35 kHz for 15 minutes as a pre-processing step; the incubation times for C1, C2, and C3 solutions were extended (30 min at 65 for C1, and 15 min at 4 for C2 and C3); bead-beating step was replaced with horizontal vortexing for 10 min at 2200 rpm (IKA VXR basic Vibrax shaker). The DNA quality and purity was examined by NanoDrop ND-1000 V3.8 spectrophotometer (Thermo Fisher). To gain information on bacterial community composition a portion of the DNA was sent for amplicon sequencing of the V4 region of the 16S rRNA gene by 515F and 806R primers (Apprill *et al*. 2015; Parada *et al*. 2016). In parallel, to gain information on diatom community composition, a portion of the same DNA was used to sequence a 312 bp barcode of the *rbcL* gene by combining the forward (DiatrbcL1F_708F_1, DiatrbcL2F_708F_1 and DiatrbcL3F_708F_1) and reverse (DiatrbcL1R_708F_1 and DiatrbcL2R_708F_1) primers from the Vasselon *et al*. (2017) study into one forward primer (5’-AGGTGAAYWAAAGGTTCWTAYTTAAA-3′) and one reverse primer (5’- CCTCTAATTTACCWACNACWG-3′) as listed in Supplementary Table S2. The sequencing was performed on the Illumina platform with MiSeq 250×2 bp paired-end chemistry at MrDNA (Texas, USA).

#### Diatom monoclonal cultures total DNA extraction and NGS sequencing

Diatom strains were maintained in culture for at least a year and passed several rounds of subculturing before harvesting the pellet for bacterial profiling (Table 1, Supplementary Table S1). The strains were then grown in duplicates in 75 cm^2^ cell culture flasks as described above and were harvested upon reaching late exponential phase. The cells were collected and pelleted in 50 ml conical sterile tubes by centrifuging at 5,000 g for 10 minutes prior to removing the supernatant. The pellet was transferred to a sterile 2 ml microtube, pelleted again by centrifuging at 16,000 g for 10 minutes with supernatant removed and then stored at −80 °C until DNA extraction. The DNA was extracted by DNeasy PowerLyzer Microbial kit (Qiagen) following the manufacturer’s instruction with several modifications: the cultures were disrupted by bead-beating with a mixture of sharp carbon and glass beads at 30 Hz, SL buffer was heated to 60 °C before use, and the elution buffer was incubated on the filter for 5 minutes before centrifugation and final elution of DNA. The quality and quantity of DNA were checked with Nanodrop and Qubit prior to sending each replicate’s DNA for sequencing of the V4 region of 16S rRNA gene with the 505F and 806R primers (Apprill *et al*. 2015; Parada *et al*. 2016) at Microsynth (Switzerland).

#### Bacterial cultivation from diatom monoclonal cultures

To survey the culturable bacteria within the diatom cultures, ten diatom cultures (Table 1) were harvested in late exponential phase and used to culture bacteria. The diatom cultures were grown as described above in 25 cm^2^ cell culture flasks (15-20 ml f/2) in duplicate and, upon reaching sufficient density, harvested in 15 ml conical sterile tubes for further processing.

To increase the chances for successful bacterial isolation, two approaches were used in diatom pellet pre-treatment before culturing bacteria: crushing and washing the pellets.

##### Crushed pellets

The 15 ml tube was centrifuged at 8,000 g for 10 minutes before removing the supernatant up to 1 ml of residual pellet and media. The 1 ml of material was transferred to a 1.5 ml sterile microtube and centrifuged at 16,000 g for 5 minutes before removing the residual supernatant. The pellet was then crushed by a sterile plastic pestle attached to an electric screwdriver for 5 sec. After crushing the pellet, the material was resuspended in 200 μl of sterile 0.9% NaCl solution and serially diluted. One hundred μl of each serial dilution (from 10^0^ to 10^−6^) was plated on Marine Agar (MA; Difco, Detroit, USA) plates and incubated at 20 °C for 48 hours, and then if growth was visible the plates were incubated at 15 °C for 96 more hours, or if the growth was not visible the plates were incubated at 20 °C for 96 more hours.

##### Washing pellets

The pellet was transferred to a round bottom sterile tube prior to washing several times with 10 ml sterile 0.9% NaCl solution (8,000 g 10 min 3 times), after which the pellet was resuspended in 1 ml 0.9% NaCl and serially diluted before plating on the MA. The incubation and growth conditions were as described above.

Single bacterial colonies were inspected under a stereo microscope, replated, and incubated for 2-6 days (until growth was visible) at 20 °C. Once pure, bacterial strains were collected from plates into Microbank™ vials (Fischer Scientific) for cryopreservation. Some colonies of the pure strains were collected using a pipette tip and resuspended into an alkaline lysis buffer for DNA extraction (Niemann *et al*. 1997).

For identification of the bacterial isolates 16S rRNA genes were amplified by PCR using pA (8f) and pH (20r) (Edwards *et al*. 1989) primers in 25 μl volume reactions containing: 2 μl of alkaline lysis DNA template, 2.5 μl dNTPs, 2.5 μl Qiagen PCR buffer 10x, 0.25 μl of 10 μM primers, 0.5 μl of Qiagen Taq DNA polymerase, and 17 μl Milli-Q water. The thermocycling conditions were: intial step 95 °C for 5 min, 3 cycles of 95 °C for 1 min, 55 °C for 2:15 min, 72 °C for 1:15 min; 30 cycles of 95 °C for 35 sec, 55 °C for 2:15 min, 72 °C for 1:25 min, and the final extension of 72 °C for 7 min. The products were inspected on agarose gel and purified using Nucleofast PCR purification plates (Macherey-Nagel, Dueren, Germa) according to manufacturer’s instructions. For initial identification, the V1-V3 region was sequenced with BLK1 primer (Cleenwerck *et al*. 2007) at Eurofins Genomics (https://eurofinsgenomics.eu/). In a second round the amplicons were completely sequenced with additional primers (Coenye *et al*. 1999). All primers used in this study are listed in Supplementary Table S3.

### Bioinformatic and data analyses

#### Diatom and bacteria marker gene sequences processing

Sequences for *rbcL* gene obtained by Sanger sequencing were inspected for quality and assembled into a contig in Geneious Prime v2022.0.2. 16S rRNA sequences were assembled and checked for quality using BioNumerics 7.6.3 (Applied Maths) and identified using EZBioCloud (Yoon *et al*. 2017; https://www.ezbiocloud.net). The sequences were aligned in AliView v1.28 (Larsson 2014) using MUSCLE (Edgar 2004). To be able to compare the diatom and bacterial cultures marker genes to NGS data, the full length *rbcL* and 16S sequences were trimmed to their corresponding NGS regions in AliView. Maximum likelihood phylogenetic trees for full length marker gene sequences were generated by IQ-TREE (using ModelFinder and UFBoot2=1000) and visualized by Interactive Tree of Life (iTOL) (Nguyen *et al*. 2015; Kalyaanamoorthy *et al*. 2017; Hoang *et al*. 2018; Letunic and Bork 2021). GenBank accession codes for all diatom and bacterial strains can be found in Supplementary Table S2.

#### Amplicon sequencing bioinformatic and statistical analyses

Source sample (carapace and skin) sequences obtained from MrDNA were pre-processed by FASTqProcessor (MrDNA) to remove all non-biological sequences (primers, linkers, adapters) prior to importing the data to QIIME 2 in “EMP protocol” multiplexed paired end fastq format. Diatom cultures sequences obtained from Microsynth were already trimmed and were imported to QIIME 2 in the Cassava 1.8 paired end demultiplexed format. Both *rbcL* and 16S data were processed with QIIME 2 v2021.4 (Bolyen *et al*. 2019), with the same tools but with specific parameters for each sequencing batch described in detail in resources provided in the Data and code availability section. The imported sequences were demultiplexed by q2-demux and denoised by q2-dada2 (DADA2; Callahan *et al*. 2016) which produced amplicon sequence variants (ASVs, 100% operating taxonomic units). Up to this point each sequencing batch was processed separately to reduce denoising errors and were merged accordingly after the DADA2 output; diatom *rbcL* source samples in one group; 16S diatom monoclonal culture sample replicates in second group, and 16S source samples data in third group (see Data and code availability). Further, analyses of 16S data were performed separately for diatom culture replicates, merged diatom replicates data, source sample data, and merged diatom and source sample data. Sequences were aligned by MAFFT (Katoh 2002) and FastTree2 in q2-phylogeny plugin was used to construct a phylogenetic tree (Price, Dehal and Arkin 2010). Taxonomy was assigned to ASVs through q2-feature-classifier (Bokulich *et al*. 2018, Robeson *et al*. 2021) classify-sklearn naive Bayes taxonomy classifier in SILVA v138 99% 505F-806R nb classifier (Pruesse *et al*. 2007) for 16S reads, and Diat.barcode v10 for diatom *rbcL* reads (Rimet *et al*. 2019). Reads assigned to chloroplasts and mitochondria were removed from the 16S data before further processing.

To investigate alpha and beta diversity the whole 16S dataset was rarefied to 32660 reads per sample based on inspection of rarefaction curves via q2-diversity plugin. Alpha diversity indices (Shannon’s entropy, Pielou’s evenness, Faith’s phylogenetic diversity, observed ASVs) were calculated via q2-diversity plugin. Beta diversity was explored via q2-diversity plugin on rarefied data with Bray-Curtis, Jaccard, unweighted Unifrac, weighted Unifrac (Lozupone and Knight 2005; Lozupone *et al*. 2011), and generalized Unifrac (Chen *et al*. 2012) distances. Unrarefied data was analysed through robust Aitchison distance via q2-deicode plugin (Gloor *et al*. 2017; Martino *et al*. 2019). Principal Coordinates Analyses (PCoA) for Bray-Curtis, Jaccard, all Unifrac distances, and Principal Components Analysis (PCA) for robust Aitchison (rPCA) were performed by q2-diversity and q2-deicode, respectively. Multi-way Permutational Multivariate Analysis of Variance i.e., Adonis PERMANOVA, (Anderson 2001) was used to estimate the relative impact of factors (turtle host ID and diatom genus) on the bacterial communities in diatom cultures (permutations=9999, {vegan} v2.5-7, Oksanen *et al*. 2020; {pairwiseAdonis} v0.4, Arbizu 2017). Data exploration and visualizations were performed with R v4.1.1 in RStudio (R Core Team 2021, {qiime2R} v0.99, Bisanz 2018; {tidyverse}, Wickham *et al*. 2019; {ggplot2}, Wickham 2016; see Data and code availability). To investigate the cultured diatom and bacterial strains presence in the NGS data, trimmed marker gene sequences were merged with ASVs, aligned in AliView, processed with IQ-TREE and visualized by iTOL as described above.

#### Data and code availability

Diatom and bacterial strains used in this study are available at the BCCM/DCG and LMG culture collections (see culture codes in Supplementary Table 2). Raw amplicon sequences (with removed non-biological sequences) are available at the European Nucleotide Archive (ENA) under the accession numbers PRJEB47668 (diatom monoclonal culture 16S rRNA sequences), PRJEB51458 (total sea turtle surface 16S rRNA sequences; sample accession numbers ERS10917111, ERS10917104, ERS10917103, ERS10917093, ERS10917093, ERS10917091), and PRJEB51297 (total sea turtle surface *rbcL* sequences). Full *rbcL* sequences per diatom strain and 16S rRNA gene sequences per bacterial isolate are available in GenBank (*rbcL* OM686876-OM686892; 16S OM959184-OM959220 and ON040652-ON040654; accession numbers per strain are listed in Supplementary Table S2). All other data supporting the conclusions in this manuscript are available in supplementary materials. Data and code used for bioinformatic analyses, statistical analyses, and data visualizations are available at GitHub (https://github.com/kl-fil/Filek_et_al._2022-diatom_microbiota) and Mendeley Data (DOI: 10.17632/4r6568xcpw.1).

**Table 2.**
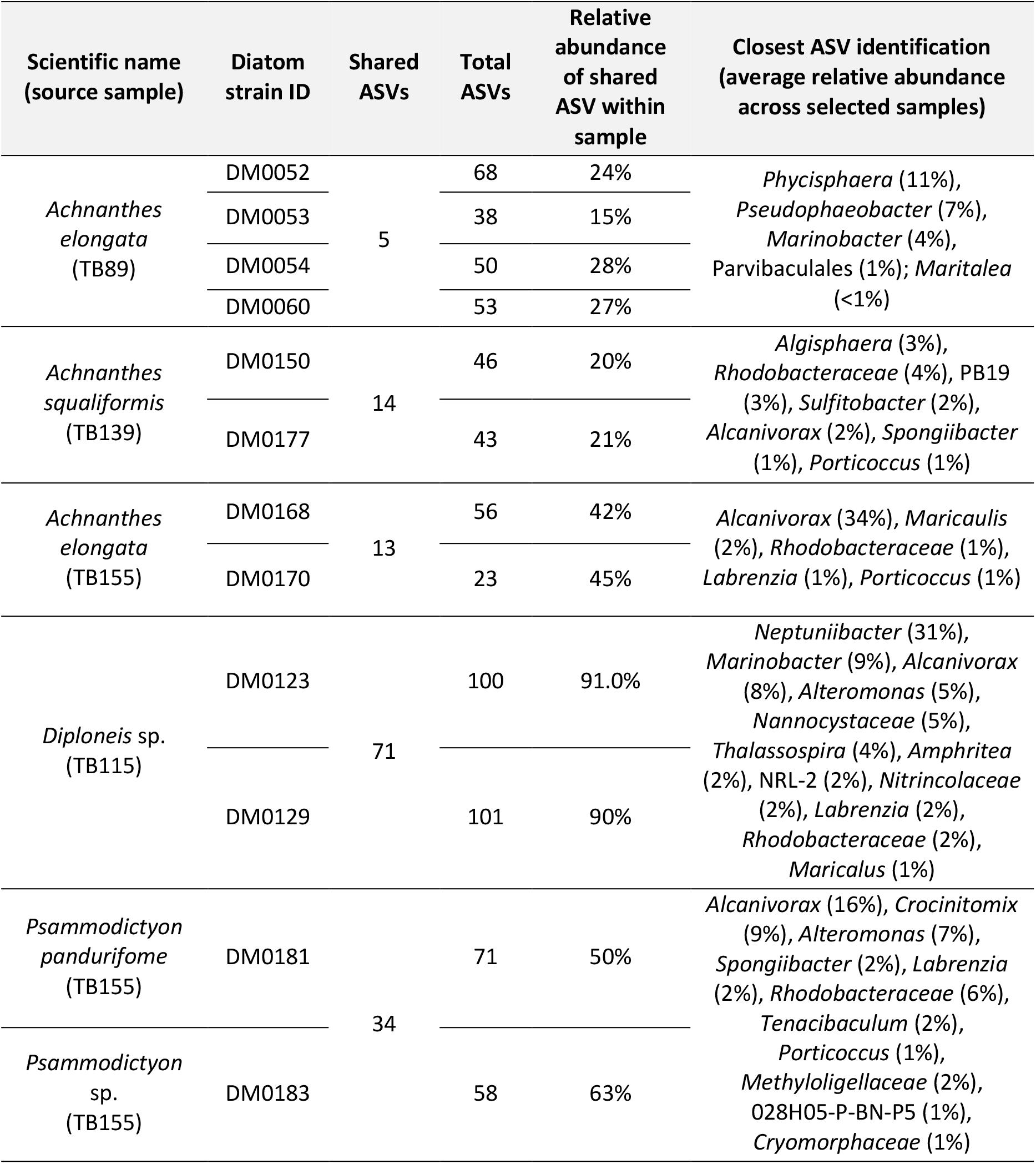
Shared ASVs within diatom genera originating from the same source sample with closest ASV identification for ASVs present above 1% average relative abundance.

## Results

### Loggerhead-associated diatom monoclonal cultures and source sample diatom community composition

Within this study we isolated and cultivated diatom strains of diverse diatom taxa and established xenic monoclonal cultures. Only cultures without detected eukaryotic contaminants were chosen for this study (Table 1). Isolated diatoms were identified as belonging to eight different genera and eleven species (Figure 1B-L), including the putative epizoic diatoms *Achnanthes elongata, Achnanthes squaliformis*, and *Poulinea lepidochelicola* (Figure 1B, C, H, M). *Achnanthes* and *Poulinea* in cultures exhibited high polysaccharide secretion in the form of stalks or mucus sheaths enabling cells to connect and form chains and/or colonies (Figure 1M), and attach to the cell culture flask surfaces. Other genera did not show such behaviour under the conditions in this study except for colony formation of *Amphora* sp. 2 (DM0182) where cells tended to cluster together in the water column and rarely attached to the cell flask surfaces. *Diploneis, Amphora, Nitzschia, Fallacia*, and *Psammodictyon* strains readily attached to surfaces, but formation of chains or stalks was not detected. Relative relationships between diatom strains in this study (except *Nitzschia* sp. DM0178 and *Diploneis* sp. DM0136 for which the *rbcL* sequences could not be obtained) based on *rbcL* marker gene are shown within the maximum likelihood phylogenetic tree in Supplementary Figure S1A.

**Figure 1.**
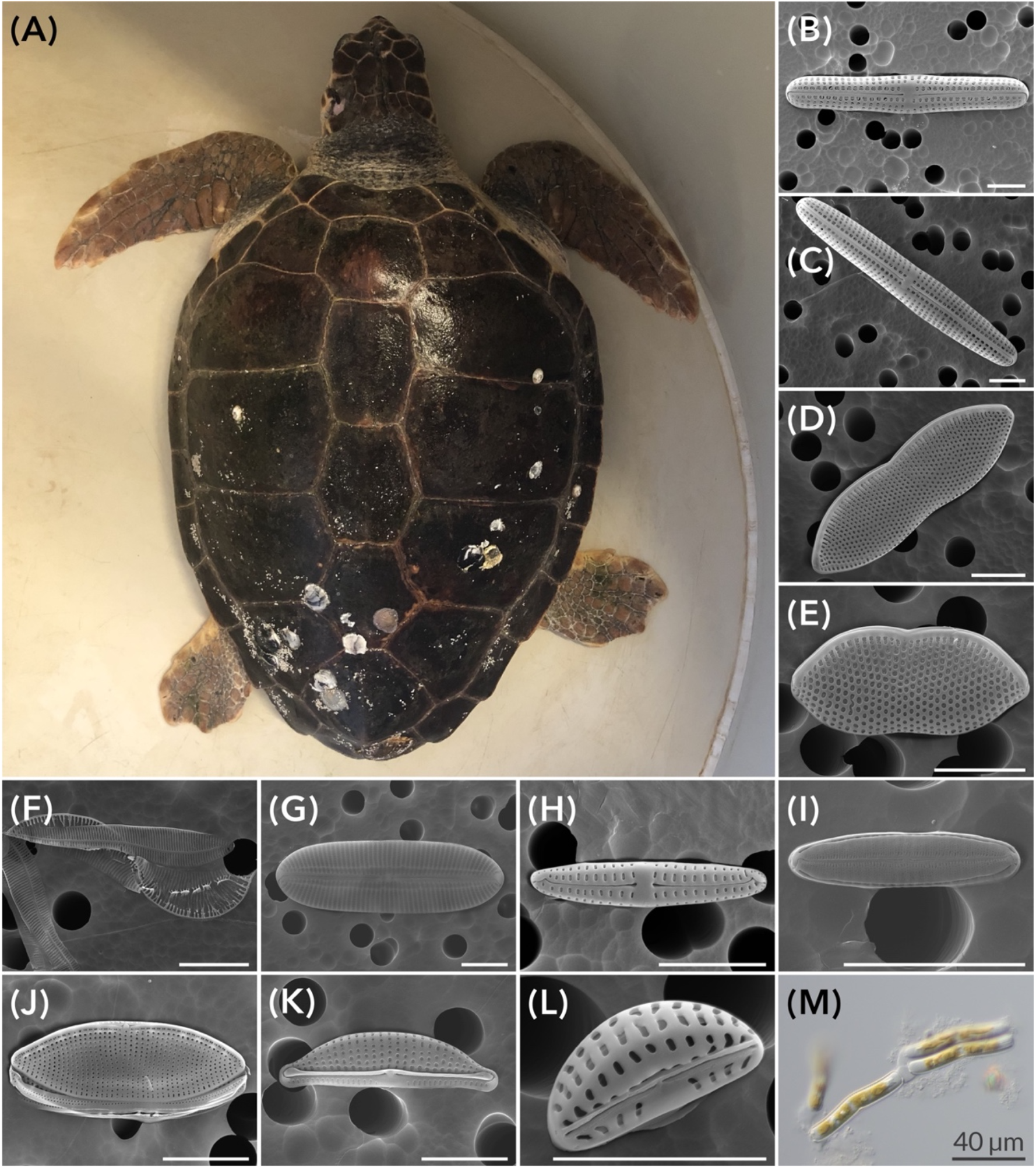
Loggerhead sea turtle and isolated diatoms. Loggerhead sea turtle Merry Fisher ID010 **(A)**, and scanning electron micrographs of diatom taxa **(B-L)**: *Achnanthes elongata* **(B)**, *Achnanthes squaliformis* **(C)**, *Psammodictyon* sp. **(D)**, *Psammodictyon panduriforme* **(E)**, *Entomoneis* sp. **(F)**, *Diploneis* sp. **(G)**, *Poulinea lepidochelicola* **(H)**, *Fallacia* sp. **(I)**, *Nitszchia* sp. **(J)**, *Amphora* sp. 1 **(K)**, *Amphora* sp. 2 **(L)**, light micrograph of *A. elongata* cells in monoculture **(M)**. All scales are 5 μm unless marked differently.

Next-generation sequencing (NGS) of the *rbcL* marker gene amplicons in source samples yielded 458,069 high quality *rbcL* sequences (median = 37,756.5) across 619 ASVs. The samples showed high proportions of *Nitzschia, Amphora, Halamphora*, and *Navicula* genera along with unclassified ASVs based on Diat.barcode taxonomy classifications of the *rbcL* marker region (Supplementary Figure S1B). However, when the NGS *rbcL* amplicon marker was extracted from full size *rbcL* sequences obtained for diatom strains, and compared to the NGS results sequence annotations, we observed discrepancies in Diat.barcode assigned taxonomy for barcodes associated with newly described epizoic taxa *A. elongata, A. squaliformis*, and *P. lepidochelicola* (see Supplementary Figure S2). Positioning of diatom strain extracted *rbcL* barcodes in the phylogenetic tree (Supplementary Figure S2) coincided with NGS sequences annotated as *Nitzschia* spp. (DM0052, DM0053, DM0054; *A. elongata*), *Bacillariaceae* (DM0060, DM0168, DM0170; *A. elongata*), *Amphora* spp. (DM0077; *P. lepidochelicola* and DM0150, DM0177 *A. squaliformis*) with low confidence values (Supplementary Table S4). Alignment of NGS *rbcL* ASVs and *rbcL* barcode sequences of diatom strains found exact matches for most diatoms except *Fallacia* sp. (DM0147), *Psammodictyon panduriforme* (DM0181), *Amphora* sp. 2 (DM0182), and *Psammodictyon* sp. (DM0183). Matched *rbcL* ASVs were present in source samples mostly around 1% relative abundance. Interestingly, *Amphora* sp. 1 (DM0070) matched ASV was present at 48% relative abundance in its source sample, while *P. lepidochelicola* (DM0077) was present at 32% in its source sample (Supplementary Table S4), thus forming a significant portion of the diatom assemblage of the turtle associated microbial biofilm. Epizoic diatoms *A. elongata* and *A. squaliformis* were present in their corresponding source samples at less than 1% and at 3% relative abundance, respectively. For other strains that could not be matched to an *rbcL* ASV we examined the closest neighbours in the phylogenetic tree (Supplementary Figure S2) and their presence was also around 1% in at least one source sample (Supplementary table S4).

### Bacterial communities of source samples (carapace and skin) and diatom monoclonal cultures

Source samples yielded 856,010 16S rRNA gene sequences (median=159,904.5) across 4,275 ASVs. Diatom cultures (19 strains in two replicates, n_(total)_ = 38) yielded 2,297,642 high quality 16S rRNA gene sequences (median = 63,314.5) across 485 ASVs. Chloroplast reads encompassed 497,842 sequencing reads in diatom cultures NGS data; on average 21% of reads across all samples were associated with chloroplasts (ranging from 1% to 75% of relative abundance). After filtering chloroplast and mitochondria sequences, source samples retained 3661 ASV and diatom monocultures 458 ASVs; with 216 ASVs in common. Shared ASVs comprised an average 40% relative abundance (SD ± 0.2) of diatom associated bacteria, and average 8% relative abundance (SD ± 0.6) of source sample bacterial community (Supplementary Table S5).

### Within sample bacterial community diversity

Source samples exhibited high alpha diversity (Supplementary figure S3) with 917 ASVs on average. Diatom cultures contained 52 ASVs on average (spanning from 18 to 101) with most other alpha indices several times lower than in source samples (Supplementary figure S3). Pielou’s evenness and Faith’s phylogenetic index showed some diatom cultures are dominated by specific ASVs and lack phylogenetic diversity, while others heave a more equal prevalence of ASVs and higher diversity. No metadata categories were found to be responsible for such observations.

### Bacterial community composition and structure

Relative abundance of microbial taxa (Figure 2) shows general reduction in number of bacterial phyla in diatom cultures versus source samples (Figure 2B). Diatom monocultures contained 17 phyla in total, reduced in comparison to source samples which contained 36 (Supplementary table S5). Source samples completely lacked Elusimicrobiota phylum, which was found in only one diatom culture (*A. elongata*, DM0052) and contained only one ASV (1606) belonging to *Elusimicrobium* genus at the relative abundance of 0.2% (159 out of 78562 reads). *Rhodobacteraceae* were abundant both in diatom and source samples, while *Thalassospiraceae* and *Stappiaceae* seem to be enriched in monocultures while being part of the rare biosphere at less than 0.1% of relative abundance (Pascoal, Costa and Magalhães 2021) in source samples (Figure 2C). In class Gammaprotebacteria the effect of enriched taxa is more pronounced as families *Alteromonadaceae, Collwelliaceae, Marinobacteaceae, Alcanivoracaceae*, and *Nitricolaceae* are more abundant in monocultures (Figure 2D). Several taxa within Bacteroidota phylum also show the enrichment pattern (*Crocinitomicaceae*, Sphingobacteriales NS11.12) (Figure 2E), but *Phycispheraceae* within Planctomycetota show higher abundance in only a subgroup of mostly epizoic diatom strains originating from TB89, TB90 and TB139 source samples, even though they are barely present in TB89 and TB90 (Figure 2F).

**Figure 2.**
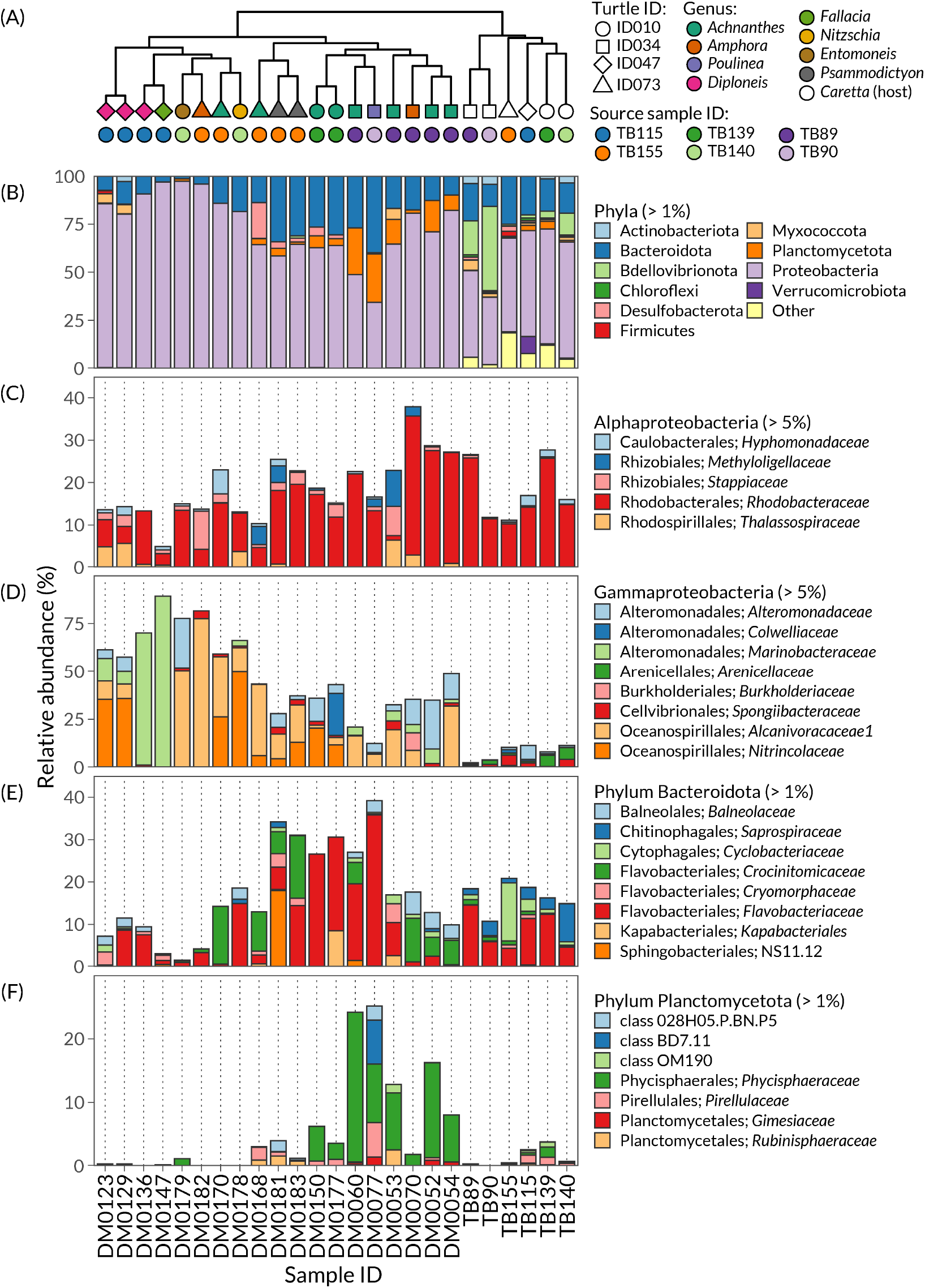
Relationships between samples and relative abundance of bacterial taxa in cultivated diatom strains and source samples. Generalized Unifrac dendrogram shows relationships between samples **(A)** with indication of diatom genus (color) and turtle ID (shape) at the tips, while specific source samples (colors) are indicated in symbols beneath. Relative abundances of bacterial taxa are presented at the levels of bacterial phyla above 1% **(B)**, classes Alphaproteobacteria **(C)** and Gammaproteobacteria **(D)** above 5%, phyla Bacteroideota **(E)** and Planctomycetota **(F)** above 1% relative abundance in at least one sample. Most common orders and families or closest taxonomic identification are shown.

### Shared bacterial taxa and individual ASVs

Source and monoculture samples shared bacterial families *Rhodobacteriaceae* and *Flavobacteriaceae*, while 85% (21/25 samples) shared additional *Hyphomonadaceae* and *Stappiaceae*. At the genus level 80% of samples (20/25) shared unclassified members of *Rhodobacteraceae, Alcanivorax* and *Labrenzia*. Source samples shared 50 ASVs of which 41 seem to be part of the rare biosphere (at less than 0.01% on average across samples) and rarely present in diatom monocultures. Only ASV 1206 (uncultured *Oligoflexaceae*) reached a relative abundance of 10% and 43% in source samples. Source samples exhibited much higher diversity than diatom cultures and consistently harboured members of Proteobacteria, Bacteriodota, Bdellovibrionota, Actinobacteriota, Myxococcota, Planctomycetota, Verrucomicrobiota, Cyanobacteria, Deinococcota, Desulfobactaerota, Chloroflexota, Firmicutes, Campilobacterota, Spirochaetota, and SAR324 Marine group B phyla (Supplementary Table S5).

All diatom strains shared Proteobacteria and Bacteroidota, while only 80% of monocultures (15/19) shared Planctomycetota phylum. *Alcanivorax* spp., unclassified *Rhodobacteraceae, Labrenzia* spp., *Marinobacter* spp., *Methylophaga* spp., and uncultured Parvibaculales are commonly found in monocultures (75% of samples). ASV 2509, classified as uncultured Parvibaculales, was found in 15/19 diatom cultures and in 2/6 source samples below 1% of relative abundance. Interestingly, diatom cultures originating from the same source shared 3-5 ASVs, except in the TB139 source group where the two cultures were *A. squaliformis* as they shared 14 ASVs. Similarly, *Achnanthes* strains originating from the source sample TB155 shared 13 ASVs (out of 45 and 43 total observed features), but the *Achnanthes* strains originating from TB89 shared only 5 ASVs. The difference between these three groups of *Achnanthes* is that source samples TB139 and TB155 had up to two to three times higher ASVs to begin with, in comparison to TB89. All *Achnanthes* strains shared just the previously mentioned ASV 2509, while *Achnanthes* from TB139 and TB155 shared 4 ASVs (belonging to *Alcanivorax* spp., *Porticoccus* spp., unclassified Parvibaculales, and *Labrenzia* spp.) (Table 2). Notably, *Diploneis* sp. strains (DM123, DM129) originating from TB115 shared 71 ASV (out of 100 and 101 total observed ASVs per strain), while *Psammodictyon* strains (DM0181, DM0183) from TB155 shared 34 ASVs (out of 71 and 58 total observed ASVs per strain) (Table 2). On the other hand, *Psammodyction* and *Achnanthes* strains from TB155 shared only 7 ASVs.

### Beta diversity analyses of bacterial communities

Taxonomic composition and individual bacterial ASV sharing between diatom strains is reflected in beta diversity metrics (Figure 3, Supplemental Figures S4 and S5). Compositional data analysis though rPCA shows a general pattern of samples separating based on diatom genus (Figure 3A) and carapace or skin source sample (Figure 3C). Highly ranked feature loadings of the rPCA overlap with previously observed taxa often found in diatom strains (genera *Alcanivorax, Neptuniibacter, Marinobacter, Alteromonas, Phycisphaera*). Generalized Unifrac considers the phylogenetic distances between ASVs and their abundance in each sample, balancing between the “weight” of highly abundant taxa (weighted Unifrac) and rare taxa (unweighted Unifrac), so the PCoA accordingly shows similarities between diatom strain samples with closely related bacterial taxa (Figure 3B, 3D) and reiterates the source sample groupings observed with robust Aitchison distance.

**Figure 3.**
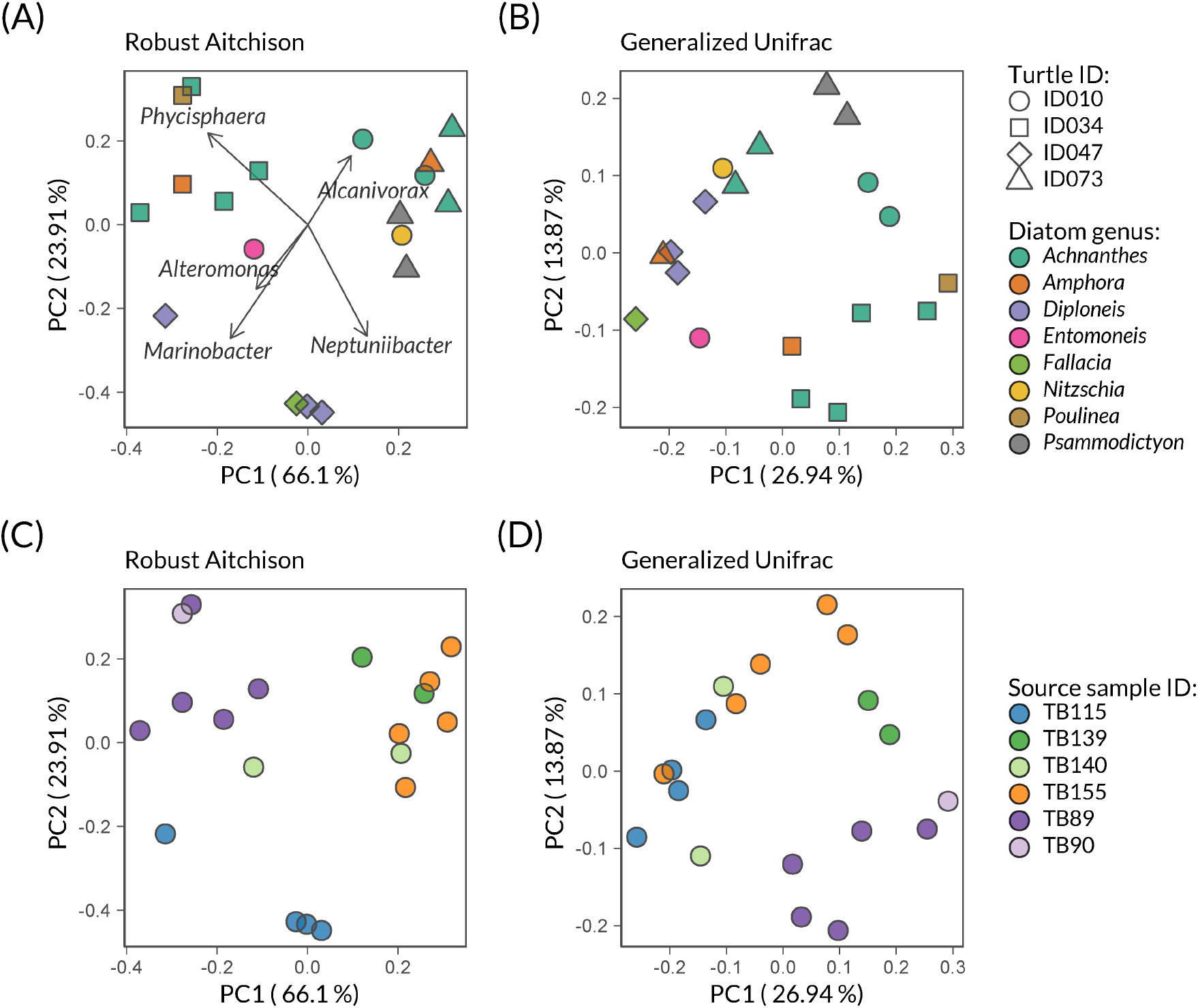
Bacterial community structure in cultivated diatom strains. Principal component analysis (PCA) of robust Aitchison distance **(A, C)** and principal coordinate analysis (PCoA) of generalized Unifrac **(B, D)** show sample clustering by turtle ID and diatom genus **(A, B)** or by specific source sample **(C, D)**.

Dominant bacterial taxa tend to drive groupings between samples based on the source sample ID while low abundance taxa tend to affect groupings in such a way that samples start reflecting the diatom genus groups (Supplementary Figures S4 and S5). Regardless, presence-absence metrics seem to separate diatom bacterial communities based on origin, revealing the environmental signature (Supplementary Figures S4C and S5C). When source samples and diatom monocultures were investigated together, despite of their extreme differences in microbial richness and diversity, above mentioned patterns recurred (Supplemental figure S6 and S7).

Multi-way Permutational Multivariate Analysis of Variance (Adonis PERMANOVA) showed significant differences when diatom monoculture samples are grouped by their individual turtle host of origin (combined carapace and skin samples; Turtle ID), and an effect of genus grouping was observed (Table 3). With Generalized Unifrac 34% of variation is explained by turtle ID (F-model=3.309, Pr(>F)=0.0001), and 35% by diatom genus (F-model=1.698, Pr(>F)=0.0007), while using the robust Aitchison 69% of variation is attributed Turtle ID (F-model=13.897, Pr(>F)=0.0001) and 15% to diatom genus (albeit genus being not significantly different in this case Pr(>F)=0.1). Pairwise ADONIS showed differences between individual turtle hosts grouped by Turtle ID ID034 vs. ID047 (generalized Unifrac F-model = 4.14, R^2^ = 0.34, Pr(> F) = 0.042; robust Aitchison F-model = 13.8, R^2^ = 0.63, Pr(> F) = 0.03), and ID034 vs. ID074 (generalized Unifrac F-model = 3.1, R^2^ = 0.25, Pr(> F) = 0.024; robust Aitchison F-model = 23, R^2^ = 0.72, Pr(> F) = 0.24).

**Table 3.** Adonis (Permanova) results based on generalized Unifrac and robust Aitchison distances for diatom strain microbial communities with two categorizations: Turtle ID as individual turtle host, and diatom genus. F statistic p-values significance level is Pr(> F) < 0.05 (in bold).

|  | Generalized Unifrac |  |  | Robust Aitchison |  |  |
| --- | --- | --- | --- | --- | --- | --- |
|  | F-model | R<sup>2</sup> | Pr(>F) | F-model | R<sup>2</sup> | Pr(>F) |
| **Turtle ID** | 3.309 | 0.34 | **0.0001** | 13.897 | 0.69 | **0.0001** |
| **Genus** | 1.698 | 0.35 | **0.0007** | 1.579 | 0.16 | 0.1 |

### Bacterial isolates from diatom monoclonal cultures

A total of 125 bacterial isolates were obtained from 10 diatom cultures (Table 1, Supplementary Table S2). Partial sequencing of the 16S rRNA gene was used to identify the strains of interest (possibly unique) and 40 strains were chosen for full 16S rRNA sequencing (Supplementary Table S2). Based on the V4 region of the full 16S rRNA sequence, 39 out of 40 bacterial strains were matched with an ASV in diatom and source sample sequencing data (Figure 4). Notably, ASVs of five bacterial isolates were not detected in the diatom strain they originated from but were detected in other diatom strains; the only bacterial strain not matched with an ASV was classified as belonging to the genus *Actibacterium* (Figure 4).

**Figure 4.**
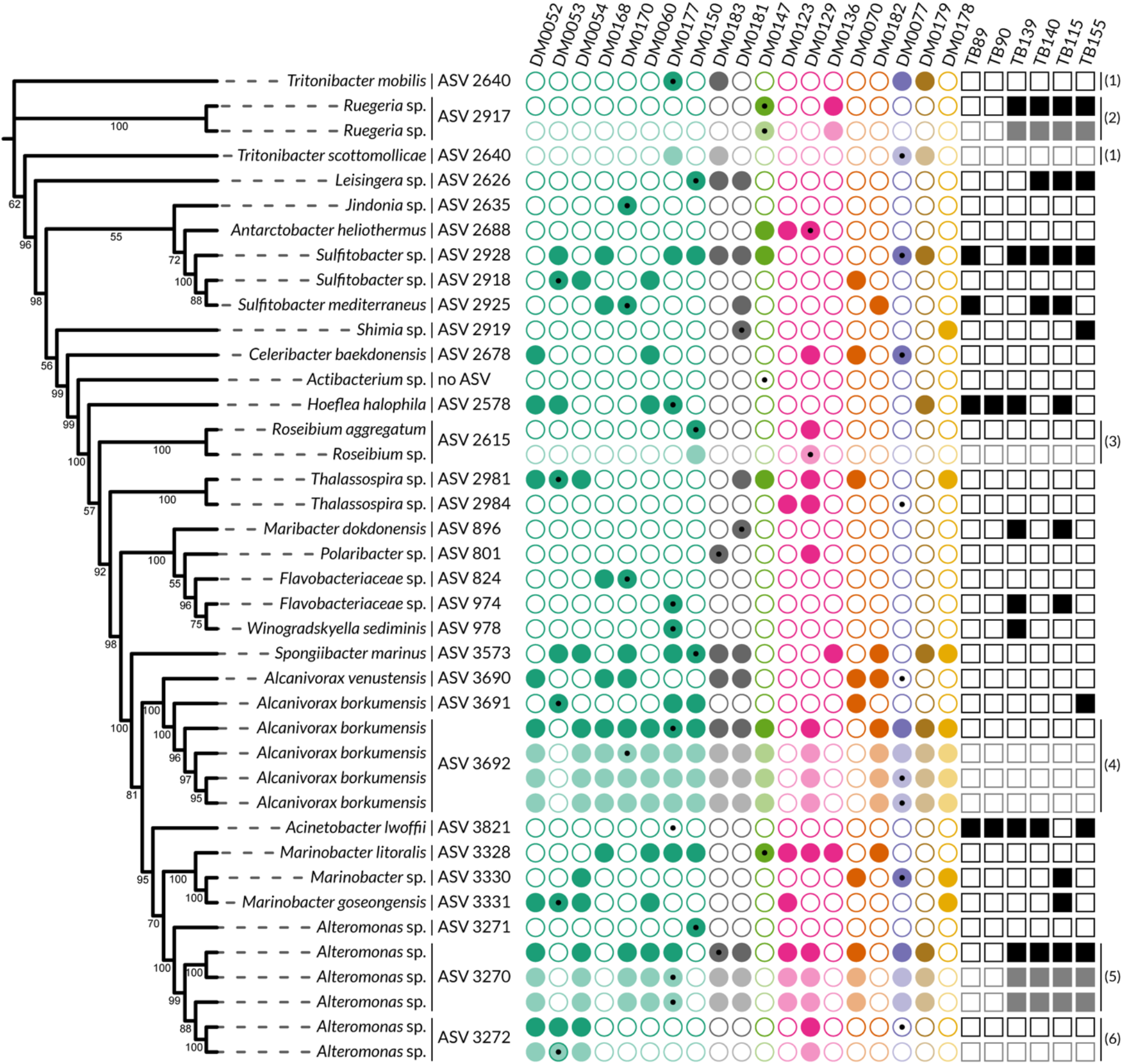
Relative relationships of bacterial isolates from diatom cultures and their matching ASV presence in diatom or source samples. Maximum likelihood phylogenetic tree of full 16S rRNA gene sequences of bacterial isolates was constructed in IQ-Tree and modified in iTOL (bootstrap values shown, branch lengths ignored). ASVs matched with bacterial isolates were identified based on the V4 region of the full 16S rDNA of bacterial isolates. Their presence or absence in the diatom and source NGS data is shown for each bacterial strain by filled or empty circles, respectively. Diatom strain of origin for each bacterial isolate is indicated by a black dot in the presence/absence matrix. As several bacterial isolates were matched with an identical ASV there is repetition in rows of presence/absence data indicated by faded colours and numbers on the side of the matrix as follows: (1) ASV 2640, (2) ASV 2917, (3) ASV 2615, (4) ASV 3692, (5) ASV 3270, (6) ASV 3272.

Based on full 16S rRNA sequence we managed to obtain identification for ASVs that were classified differently by SILVA or had assigned taxonomy only above genus level: several ASVs that were assigned to *Rhodobacteraceae* by SILVA were identified as belonging to genera *Leisingera, Jindonia, Tritonibacter, Celeribacter*, and *Antarctobacter*. ASV 2615 and ASV 2918 were assigned to genus *Labrenzia* and *Sedimentitalea* by SILVA (confidence at 0.91 and 0.72), but we identified them as belonging to *Roseibium* and *Sulfitobacter* genus, respectively. Additionally, with SILVA two ASVs (824 and 974) were assigned to genera *Aquibacter* and *Winogradskyella*, while full 16S indicates they could potentially belong to a new genus in the *Flavobacteriaceae* family. Multiple bacterial strains matched with one ASV even though they differed in full length 16S, such as *Tritonibacter scottomolliaceae* and *Tritonibacter mobilis, Alteromonas* spp., and *Alcanivorax* spp. (Figure 4, Supplementary Tables S5 and S6).

ASVs matched to bacterial isolates were detected in at least one diatom monoculture (except ASV 3821); however, only 14 out of 30 matched ASVs were present in source samples and at less than 1% relative abundance (Figure 4, Supplementary Table S6).

Interestingly, based on the ASVs, *Alcanivorax* spp. were enriched in most diatom cultures, while in source samples they were detected at less than 1% relative abundance (2-15 reads in four out of six source samples), and ASV 3821 (matching to *Acinetobacter lwofii*) was detected only in source samples and not in monocultures (Figure 4, Supplementary Table S6).

## Discussion

Investigations of diatom-bacteria associations, although crucial for understanding global ecological processes, are still limited mostly to habitats such as sediment biofilms or planktonic communities. In this study we provide a multi-level inventory of diatoms and bacteria associated with loggerhead sea turtles. Our approach combined PCR-based surveys of microbial communities (*rbcL* and 16S rRNA gene amplicon sequencing) in source samples as well as isolating and culturing non-model and novel diatom taxa, thus showing that marine reptiles are valuable “hot spots” of diatom and bacterial diversity (Hooper *et al*. 2019; Keller *et al*. 2021). As diatoms are hosts to bacteria within their phycosphere we further surveyed the bacterial community retained in diatom monoclonal cultures that revealed instances of rare-in-source bacterial taxa enrichment and potentially important diatom-bacteria associations.

### Sea turtle carapace and skin harbour diverse microbial communities

Sea turtle harbour diverse macro- and microorganisms on their carapace and skin (Frick and Pfaller 2013; Rivera *et al*. 2018; van de Vijver *et al*. 2020, Kanjer *et al*. 2022 preprint). Bacterial communities associated with sea turtle skin and carapace are rich, highly diverse, and reflect the sampling locality of the turtle (expanded in Kanjer *et al*. 2022 preprint). The carapace and skin samples in this study were obtained from turtles that were sampled before admission to rehabilitation, during rehabilitation, and post-rehabilitation after spending time in an open pool with recirculating sea water (see Supplementary Table S1) which could affect the epizoic biofilm composition. Although our study focused on a limited number of turtles, our isolation and cultivation efforts led to establishing cultures of several newly described diatom species *A. elongata, A. squaliformis, P. lepidochelicola*, and potential new species such as *Diploneis* sp., *Fallacia* sp., *Amphora* spp., *Psammodictyon* sp. (Majewska *et al*. 2015, 2017).

### Phycosphere bacterial community composition reflects diatoms’ source environment and genus

Similar to other benthic diatom-bacteria biofilms, diatoms on sea turtles can be observed in dense assemblages, surrounded by extracellular polysaccharides and bacteria either on the diatom cells or surrounding organic matter (Bosak unpublished data). Since diatom cells in this study were washed through several series of sterile growth medium during isolation, we assume that the bacteria transferred with the diatoms are the ones found in close association with the diatom cells’ phycosphere in their natural habitat therefore allowing us to reveal potentially relevant diatom-bacteria associations. Specific ASVs detected both in source samples and in diatom cultures comprised only a small proportion of total microbial community in source samples, while they made up almost half of the community in diatom cultures. Bacterial ASVs that were not detected in source samples but are present in diatom cultures could have been a part of rare taxa in source samples and possibly only became detectable once enriched in diatom cultures. Beta diversity analyses that do not consider the bacterial ASVs’ phylogenetic relationships (Jaccard or Bray-Curtis) consistently grouped the diatom strain bacterial communities based on the source sample.

However, once phylogenetic distance between bacterial ASVs was considered (unweighted, weighted, and generalized Unifrac) the strength of the source sample effect lessened, and possible effects of the diatom host “selection” were accentuated. Our results support the findings that closely related diatom species recruit bacteria from their immediate environment and retain the environmental signature in culture while also selecting for related bacterial taxa, depending on their lifestyle and functions provided by the bacterial community (Amin, Parker and Armbrust 2012; Seymour *et al*. 2017; Behringer *et al*. 2018; Crenn, Duffieux and Jeanthon 2018; Majzoub *et al*. 2019; Stock *et al*. 2019, 2022; Mönnich *et al*. 2020; Filho *et al*. 2021).

### Diatoms enrich bacterial taxa that are otherwise scarce

Studies so far have shown diatom phycosphere is usually dominated by Proteobacteria (mainly Alphaproteobacteria) and Bacteroidota phyla members: *Sulfitobacter, Roseobacter, Ruegeria, Marinobacter, Alteromonas*, and *Flavobacterium* (Amin, Parker and Armbrust 2012, 2012; Goecke *et al*. 2013; Seymour *et al*. 2017; Majzoub *et al*. 2019), and that bacterial consortia are stable over time in xenic diatom monoclonal cultures (Behringer *et al*. 2018; Barreto Filho *et al*. 2021). While we consistently observed Alphaproteobacteria in diatom cultures, we also detected a strong enrichment of members of Gammaproteobacteria, namely *Nitrincolaceae* that are usually detected in diatom blooms (Liu *et al*. 2020) and *Alcanivoracaceae* that are predominant in oil contaminated sea water (Kasai *et al*. 2002; Bookstaver *et al*. 2015). Historically, the genus *Alcanivorax* has been associated with hydrocarbon degradation, while recent studies show *A. borkumensis* is common in the plastisphere in the Mediterranean Sea with ability to degrade low density polyethylene (Delacuvellerie *et al*. 2019). *Alcanivorax venustensis* and *A. borkumensis* were readily isolated from most diatom cultures in this study as it is possible that the diatom hosts produce organic nutrients beneficial to *Alcanivorax*. To the authors’ knowledge, *Alcanivorax* genus has not yet been reported in other diatoms in culture. Even so, *Alcanivorax* has been reported in the phycosphere of dinoflagellates (Denaro *et al*. 2021), it was isolated from commercial *Nannochloropsis* cultures grown in plastic bags of ProviAPT reactors (Giraldo *et al*. 2019), and was found to be a major constituent of tidal biofilms where it could consume diatom produced hydrocarbons (Coulon *et al*. 2012).

On the other hand, Planctomycetota phylum *Phycisphaeraceae* members were enriched in only a subset of diatom strains (genera *Achnanthes* and *Poulinea*). Diatom *P. lepidochelicola* harboured uncultured Planctomycetota 028H05-P-BN-P5 and BD7-11, uncultured *Phycisphaera* sp., and genera *Blastopirellula* and *Gimesia* which have been described previously as intimately associated with macroalgal surfaces (Lage and Bondoso 2014; Bondoso *et al*. 2017; Wiegand *et al*. 2020). Members of phylum Planctomycetota are known for their uncommon traits: endomembranes, anammoxomes, reproduction by budding, and good attachment abilities (Lage and Bondoso 2014 and references therein) that might have facilitated colonization or even just close association with diatoms exhibiting high polysaccharide production during colony, chain, and stalk formation (as observed in *Achnanthes* and *Poulinea* cultures).

Bacterial isolates from diatom cultures revealed that several bacterial taxa detected by NGS were readily cultured on MA, even demonstrating the potential for new species discovery as several bacterial isolates were recognized as potential novel genera within the *Flavobacteriaceae* family. These results complement previous efforts in characterizing bacteria associated with diatom cultures (Goecke *et al*. 2013; Stock *et al*. 2019). However, our culturing efforts were limited to 10 out of 19 diatom strains used in this study and biased both by diatom strain selection and growth media selection. Ideally, expanding culturing efforts to other diatoms from sea turtle surface could provide greater diversity of culturable bacteria as observed in a recently published inventory of bacterial isolates from corals and skin of cetaceans (Keller *et al*. 2021).

### Potential factors responsible for shaping diatom associated bacterial communities

Investigations focused on laboratory cultures cannot grasp the complexities of microbial networks found in nature as bacterial communities of cultures often differ strongly from those of single cells (Crenn, Duffieux and Jeanthon 2018; Boscaro *et al*. 2022). It is rarely expected that the epizoic benthic diatoms live as single cells during most of their lifetime as they are found in biofilms. Also, we cannot assume diatom biofilms on sea turtles are limited to a single species but are often mixed. Consequently, there will be several layers and modes of interactions between diatoms, bacteria, and even other microorganisms inhabiting biofilms on sea turtles.

Importance of host anatomy and spatial structure of microbial communities is recognised in vertebrate microbiome, emphasising the effects of spatiotemporal microbiome variability from skin or gastrointestinal tract down to individual skin pores or even crypts of Lieberkühn (Conwill *et al*. 2022 and references therein). Similar efforts to investigate anatomical sites and their specific microbial communities are often undertaken in echinoderms (Jackson *et al*. 2018), marine sponges (Hentschel *et al*. 2012; Verhoeven, Kavanagh and Dufour 2017), and corals (van Oppen and Blackall 2019; Keller *et al*. 2021). In this study there are several levels of host anatomy: sea turtle carapace and skin, and diatoms either as members of complex biofilms on the turtle or as laboratory cultures. The loggerhead sea turtle carapace is a hard bone shell covered by a living epidermis with a thick outer layer of keratin scutes that are shed periodically. The macro- (position on the carapace) and microanatomy (carapace scutes’ morphology) could affect both diatom colonization, localization, and their associated bacteria composition through light and nutrient availability, probability of mechanical removal via sea currents or turtle behaviour, and colonization by other turtle epibionts (Blasi *et al*. 2021). In our study we collected the total microbial community from the carapace surface or skin and as a result could not test macro- and microanatomy effects on associated diatoms and bacteria. Additionally, at the phycosphere level, diatom colony or cell morphology could potentially affect the microbes that are in the near vicinity or attached to diatom cells directly. Naturally, large diatoms have greater cell surface that could facilitate bacterial attachment in comparison to small diatoms that could prove difficult to colonize individually (Amin, Parker and Armbrust 2012; Seymour *et al*. 2017); which could provide a possible explanation for the lowest number of observed ASVs in *Amphora* sp. 2 and *Nitzschia* sp. (around five and ten micrometres long, respectively), but not in DM0170 *A. elongata* strain (around 35-40 micrometres long). Growth in colonies could further enable diatom-bacterial associations through extensive diatom polysaccharide production, cell-to-cell attachment, branching with mucilage pad junctions, and stalk formation that could fortuitously provide extra surface and different “microanatomical” niches for closely associated bacteria.

It is already known that benthic diatoms have an extensive number of genes involved in interacting with or responding to bacteria (Osuna-Cruz *et al*. 2020), and shifts in gene expression are observed in diatom-bacteria co-cultures (Amin *et al*. 2015; Durham *et al*. 2017; Cirri *et al*. 2019). Bacterial consortia affect the productivity of several common benthic diatoms (Koedooder *et al*. 2019), while *Phaeobacter inhibens* was observed to influence bacterial community assembly in axenic diatom cultures inoculated with bacteria derived from natural sea water (Majzoub *et al*. 2019). It has also been observed that bacterial exudates can indirectly affect the sexual reproduction of *Seminavis robusta* diatom through altering the diatom’s amino acid biosynthesis and stress response (Cirri, Vyverman and Pohnert 2018; Cirri *et al*. 2019), and that co-culturing of bacteria and diatoms can inhibit diatom cell division (van Tol, Amin and Armbrust 2017) or enhance growth (Amin *et al*. 2015). Recent findings also show that bacterial symbionts use quorum sensing to promote biofilm formation and phycosphere colonization, while diatoms can directly modulate bacterial colonization and suppress the opportunist attachment via secondary metabolites such as azelaic acid and rosmarinic acid (a quorum sensing mimic) (Fei *et al*. 2020; Shibl *et al*. 2020).

Exact mechanisms of microbial community assembly in diatom hosts were not the focus of this study, but it seems diatom strains investigated here are open to diverse colonization rather than being restricted to a small number of bacterial symbiont taxa. Hosting a repertoire of diverse bacteria (and in turn harbouring their metabolic potential) could possibly prove beneficial to complex microbial communities in dynamic environments (Henry *et al*. 2021; Pfister *et al*. 2022). However, it is difficult to infer the functions and potential benefits, or lack thereof, in microbial assemblages solely through marker gene amplicon sequencing data as marker gene sequences do not reflect the ecotypes and genomic diversity in microbes (Sjöqvist *et al*. 2021). Similarly, even though in our study most diatom strain *rbcL* sequences were matched to *rbcL* ASVs in source samples, taxonomic assignment of matched *rbcL* ASVs in source samples via Diat.barcode database did not parallel our morphology-based identification of novel diatom species (e.g., genera *Achnanthes, Poulinea*, and *Fallacia*; see Supplementary Figure S1). Current *rbcL* databases still lack sequenced representatives of diatom groups such as benthic pennate diatoms, which leads to inconsistencies between sequencing and morphology data (Rivera *et al*. 2018). To investigate lesser-known diatom taxa (as in this study) and their communities, expanding diatom isolation, cultivation, and marker gene sequencing efforts is necessary. Hence, comprehensive diatom and bacteria community, and subsequently strain characterization is needed to gain insight into functions and metabolic exchanges within epizoic diatom-bacteria communities. Even though studying complex diatom biofilms remains challenging, especially on marine vertebrate hosts, a combination of culture based and culture independent approaches focusing on individual diatom taxa and their associated bacteria can provide a baseline for discovering diatom-bacteria association patterns and hypothesis generation.

## Ethics declarations

Sampling was performed in accordance with the 1975 Declaration of Helsinki, as revised in 2013 and the applicable national laws. The sampling at the Sea Turtle Clinic (Bari, Italy) was conducted with the permission of the Department of Veterinary Medicine Animal Ethic Committee (Authorization # 4/19), while sampling in Croatia was done in accordance with the authorization of the Marine Turtle Rescue Center by the Ministry of Environment and Energy of the Republic of Croatia.

## Author contributions

KF and SB, PC, WV, AW conceptualized the study. KF, LL, MŽ performed the laboratory work. Data curation, formal analysis, data visualisation, and writing of the original draft was performed by KF. SB, AW, and KF provided funding for the research. All authors were involved in revision and editing of the manuscript.

## Acknowledgments

We are grateful to Adrianna Trotta of the Sea Turtle Clinic (Bari, Italy) and Aquarium Pula (Croatia) for assistance with sample collection. We want to thank Maja Mucko, Antonija Matek, and Lucija Kanjer for their assistance in diatom strains isolation and cultivation; Martina Galeković, Olga Chepurnova, and Tine Verstraete for support with laboratory procedures.

