## Supplementary for "More than just hitchhikers: a survey of bacterial communities associated with diatoms originating from marine reptiles": Filek2022-Supplementary_data_description.docx

Supplementary data for the manuscript
Filek *et al.* 2022 More than just hitchhikers:

a survey of bacterial communities associated with diatoms originating from marine reptiles

**Supplementary figures**

**Supplementary Figure S1.** Relative relationships between diatom strains in this study and diatom community composition in source samples based on *rbcL* gene NGS sequencing. Maximum likelihood phylogenetic tree of diatom strains in this study based on full *rbcL* gene (A) was generated with IQ-tree and iTOL, and shows ultrafast bootstrap support. Relative abundance of diatom taxa (above 2% relative abundance) in source samples was obtained by amplicon sequencing of the *rbcL* gene 263 bp barcode (B).

**Supplementary Figure S2.** Maximum likelihood phylogenetic tree (generated with IQ-tree and iTOL) of NGS data of *rbcL* barcode sequences from source samples and *rbcL* sequences obtained from diatom cultures after trimming to match the *rbcL* barcode. Each diatom strain was annotated with its scientific name and corresponding symbols used throughout the manuscript (turtle ID shapes and diatom genus colors).

**Supplementary Figure S3.** Alpha diversity metrics (observed ASVs, Shannon’s entropy, Pielou’s evenness, Faith’s phylogenetic diversity) for diatom strains and source samples arranged in the same sequence as in the main manuscript (based on generalized Unifrac distance clustering). Shapes encode turtle ID while colors encode the sample organism’s genus.

**Supplementary Figure S4.** Bacterial community structure in diatom strains. Principal coordinate analyses (PCoA) of Bray-Curtis dissimilarity (A), weighted Unifrac (B), Jaccard similarity (C), unweighted Unifrac (D) show sample clustering by turtle ID (shape) and diatom genus (color).

**Supplementary Figure S5.** Bacterial community structure in diatom strains. Principal coordinate analyses (PCoA) of Bray-Curtis dissimilarity (A), weighted Unifrac (B), Jaccard similarity (C), unweighted Unifrac (D) show sample clustering by source sample ID (color).

**Supplementary Figure S6.** Bacterial community structure in diatom strains and source samples. Principal coordinate analyses (PCoA) of Bray-Curtis dissimilarity (A), weighted Unifrac (B), Jaccard similarity (C), unweighted Unifrac (D), generalized Unifrac (F) and principal component analysis (PCA) of robust Aitchison distance show sample clustering by turtle ID (shape) and diatom genus (color).

**Supplementary Figure S7.** Bacterial community structure in diatom strains and source samples. Principal coordinate analyses (PCoA) of Bray-Curtis dissimilarity (A), weighted Unifrac (B), Jaccard similarity (C), unweighted Unifrac (D), generalized Unifrac (F) and principal component analysis (PCA) of robust Aitchison distance show sample clustering by sample type (monoculture or source; shapes) and source sample ID (color).

**Supplementary tables**

**Supplementary table S1.** Extended metadata for diatom monoclonal cultures replicates and source samples sent for NGS sequencing of the V4 region of the 16S rRNA gene. The table contains information on the turtle condition during sampling, origin of the turtle, dates of sample collection, diatom isolation, diatom culture harvesting for DNA isolation, and other information relevant for bioinformatic analyses reproducibility.

**Supplementary table S2.** Diatom monoclonal cultures and bacterial strain list with culture collection codes and accession numbers for *rbcL* and 16S rRNA gene sequences, respectively.

**Supplementary table S3.** List of primers and sources used for marker gene (*rbcL*, 16S rRNA gene) Sanger sequencing for diatom or bacterial isolates, and primers used for NGS sequencing of diatom monoclonal cultures and source samples.

**Supplementary table S4.** Source samples *rbcL* gene amplicon sequencing reads and their taxonomic classification. *rbcL* barcode from diatom monoclonal cultures was extracted and matching or closest neighbour ASVs are indicated.

**Supplementary table S5.** Diatom monoclonal cultures and source samples 16S rRNA gene amplicon sequencing reads (after filtering out chloroplast and mitochondria sequences) and their taxonomic classification. V4 region barcode from the bacterial isolated was extracted and matching ASVs are indicated.

**Supplementary table S6.** Extracted matched ASVs of bacterial isolates and relative abundances of ASVs per diatom monoclonal culture and source sample.
