## Supplementary figures and images for "More than just hitchhikers: a survey of bacterial communities associated with diatoms originating from marine reptiles"

### Supplementary_Figure_S1.pdf

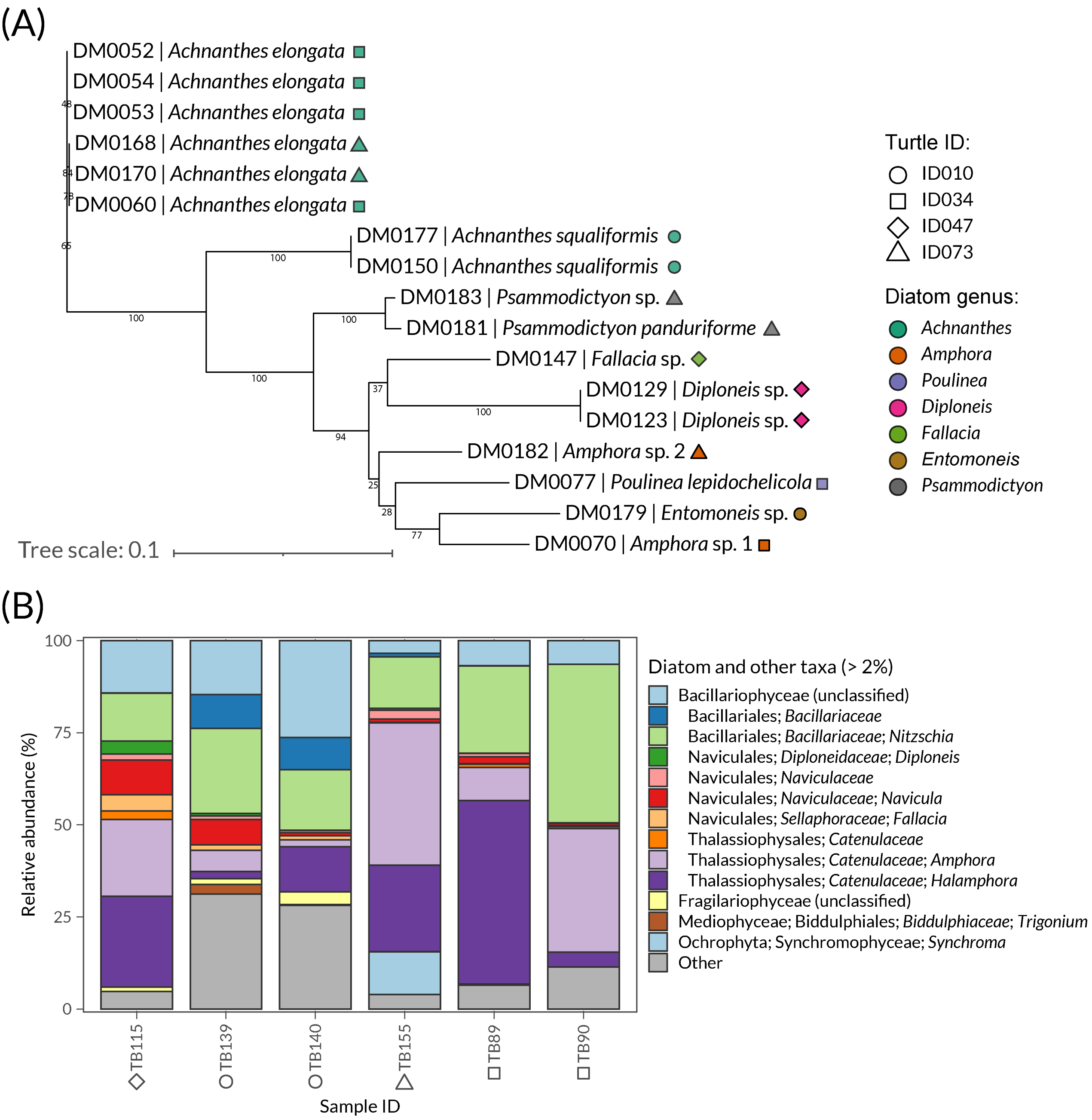

### Supplementary_Figure_S2.pdf

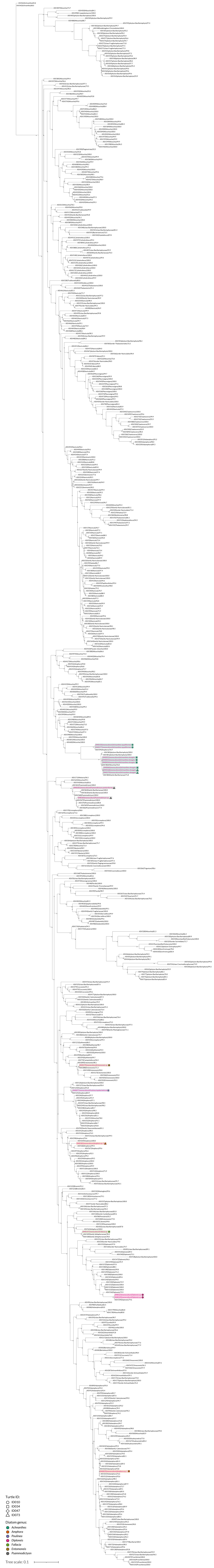

### Supplementary_Figure_S3.pdf

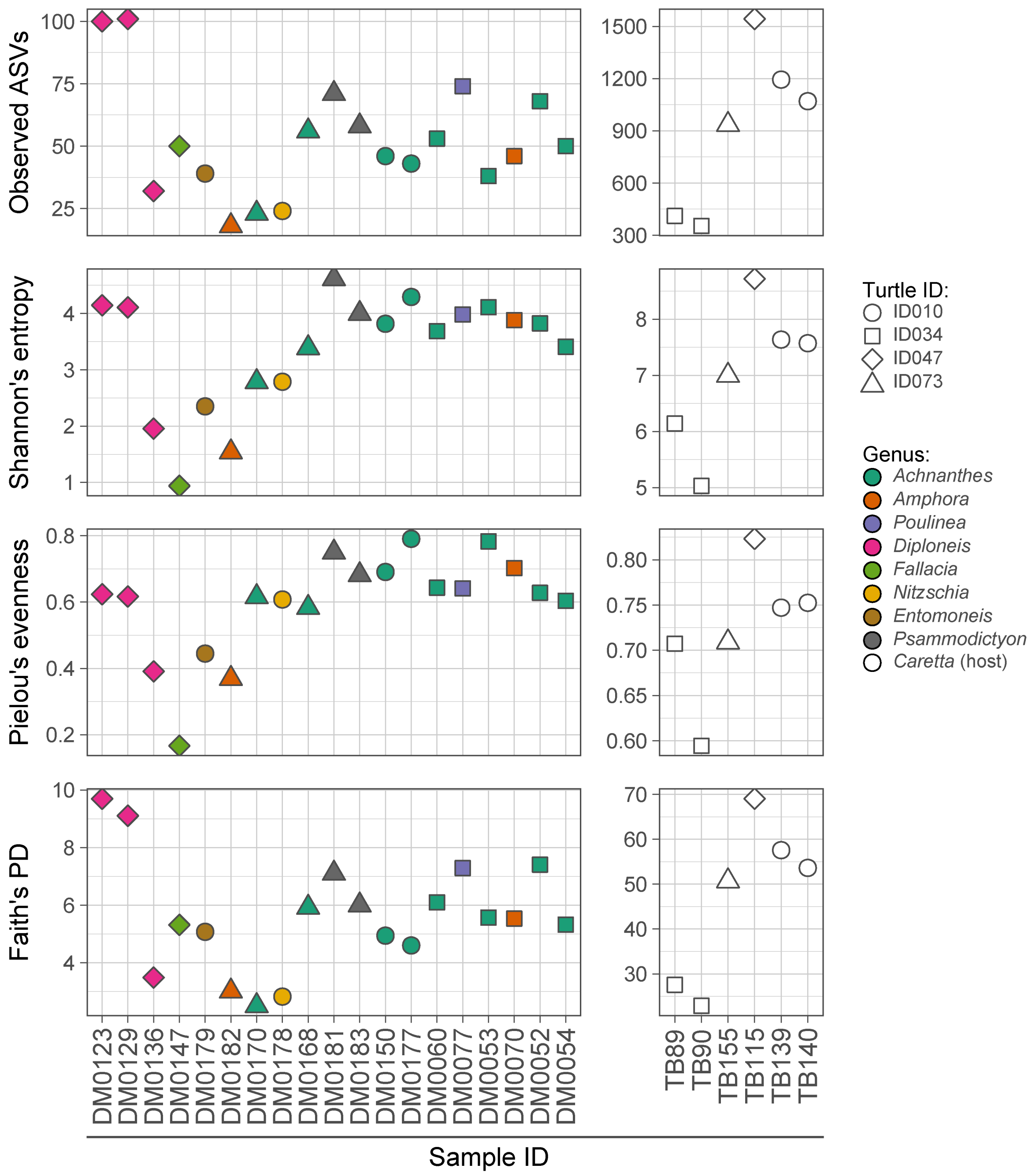

### Supplementary_Figure_S4.pdf

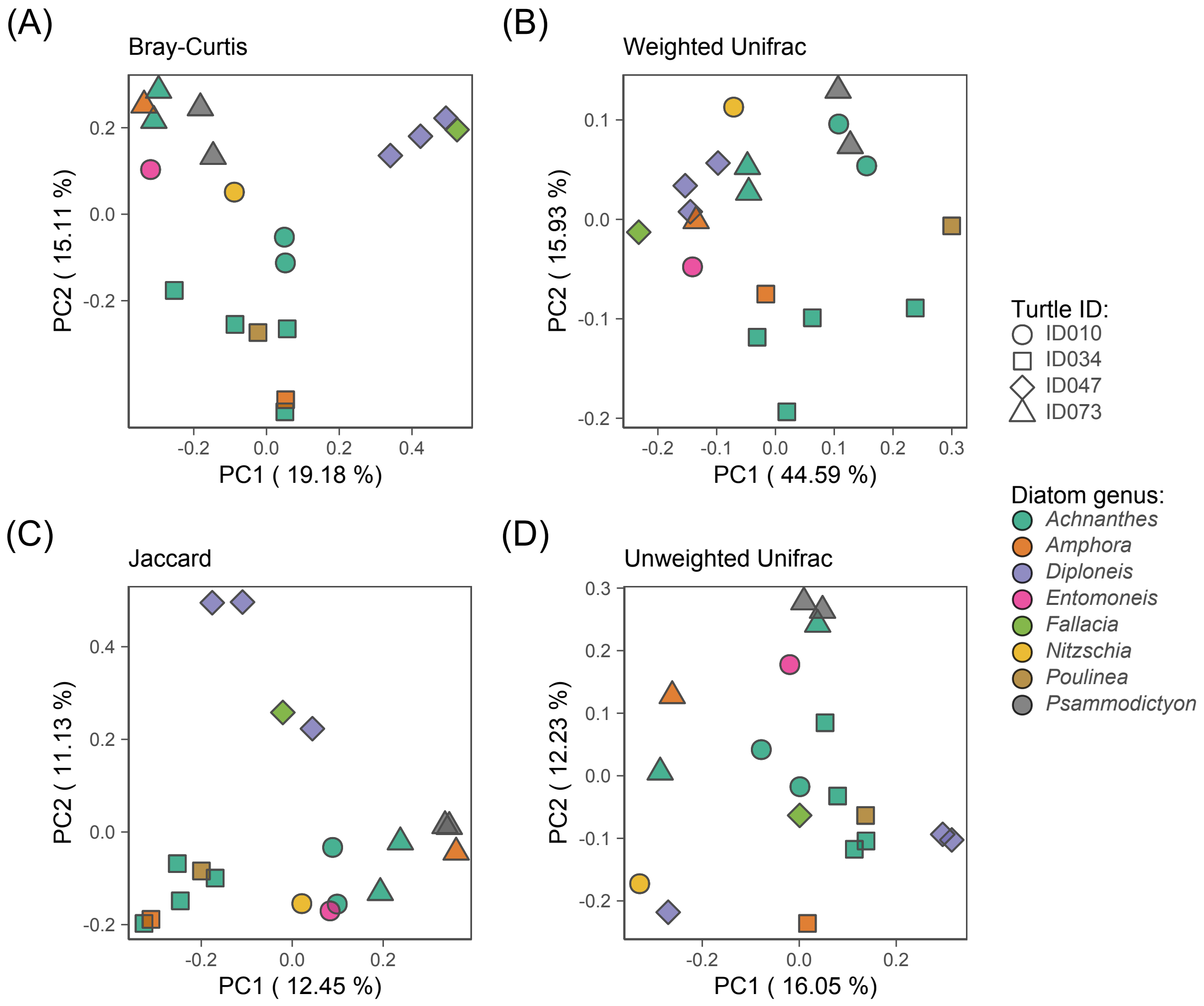

### Supplementary_Figure_S5.pdf

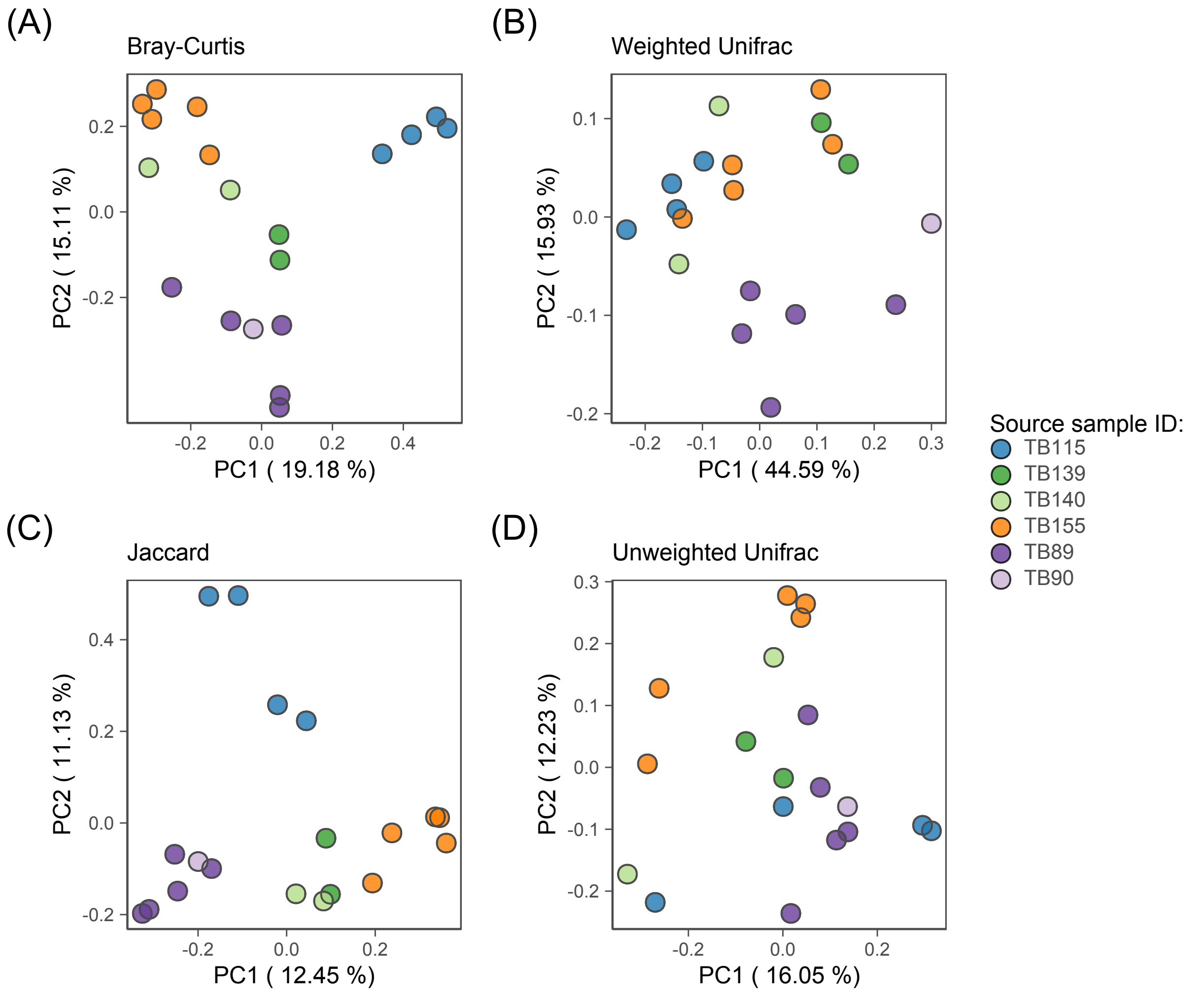

### Supplementary_Figure_S6.pdf

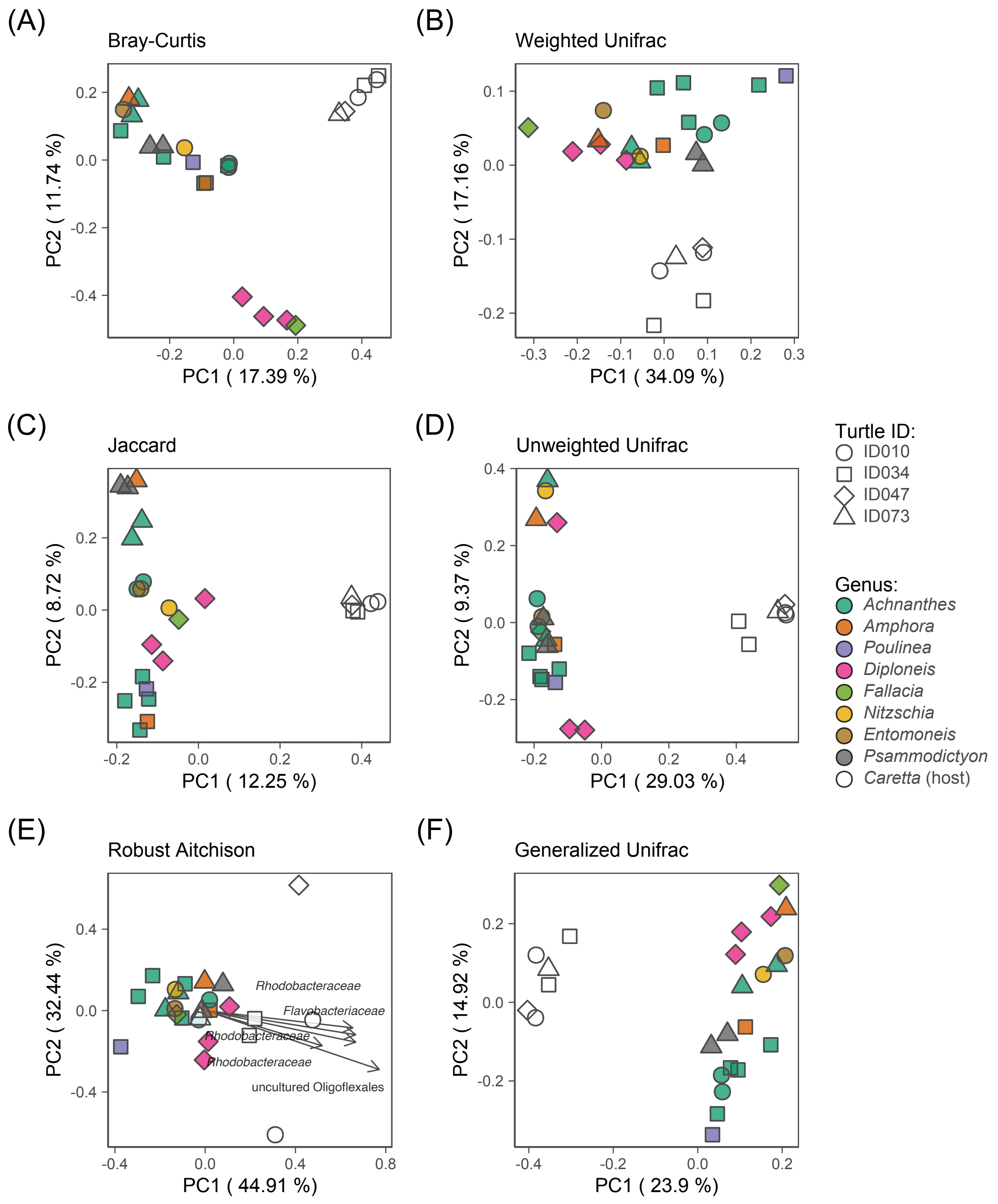

### Supplementary_Figure_S7.pdf

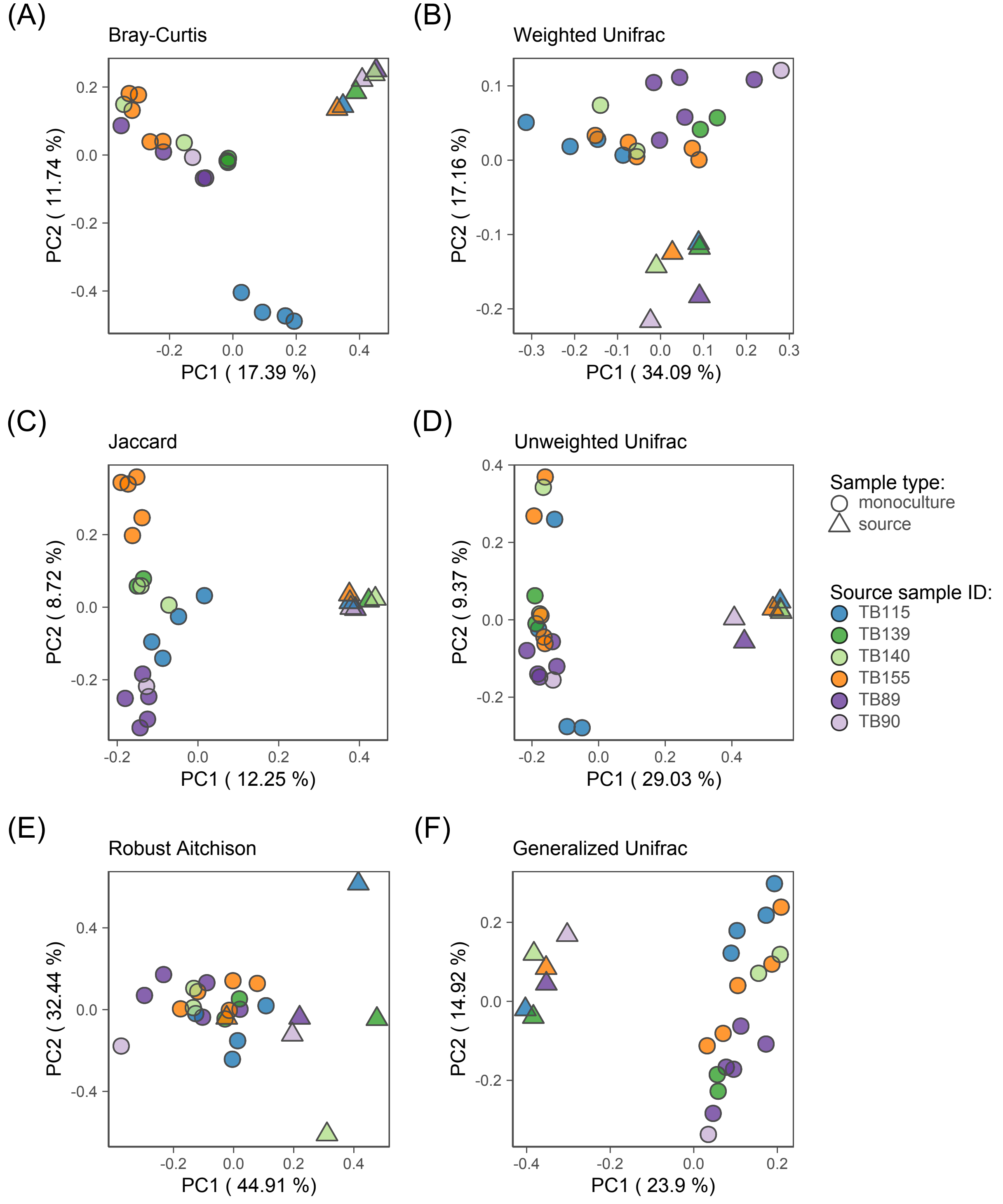
